# Aberrant accumulation of age- and disease-associated factors following neural probe implantation in a mouse model of Alzheimer’s disease

**DOI:** 10.1101/2023.02.11.528131

**Authors:** Steven M. Wellman, Olivia A. Coyne, Madeline M. Douglas, Takashi D.Y. Kozai

## Abstract

Electrical stimulation has had a profound impact on our current understanding of nervous system physiology and provided viable clinical options for addressing neurological dysfunction within the brain. Unfortunately, the brain’s immune suppression of indwelling microelectrodes currently presents a major roadblock in the long-term application of neural recording and stimulating devices. In some ways, brain trauma induced by penetrating microelectrodes produces similar neuropathology as debilitating brain diseases, such as Alzheimer’s disease (AD), while also suffering from end-stage neuron loss and tissue degeneration. To understand whether there may be any parallel mechanisms at play between brain injury from chronic microelectrode implantation and those of neurodegenerative disorder, we used two-photon microscopy to visualize the accumulation, if any, of age- and disease-associated factors around chronically implanted electrodes in both young and aged mouse models of AD. With this approach, we determined that electrode injury leads to aberrant accumulation of lipofuscin, an age-related pigment, in wild-type and AD mice alike. Furthermore, we reveal that chronic microelectrode implantation reduces the growth of pre-existing amyloid plaques while simultaneously elevating amyloid burden at the electrode-tissue interface. Lastly, we uncover novel spatial and temporal patterns of glial reactivity, axonal and myelin pathology, and neurodegeneration related to neurodegenerative disease around chronically implanted microelectrodes. This study offers multiple novel perspectives on the possible neurodegenerative mechanisms afflicting chronic brain implants, spurring new potential avenues of neuroscience investigation and design of more targeted therapies for improving neural device biocompatibility and treatment of degenerative brain disease.

## 2.0. INTRODUCTION

Penetrating microelectrodes offer investigators and clinicians a multi-modal approach for recording discrete signals from the brain and exogenously manipulating neuronal activity with high spatial and temporal resolution through stimulation^1^. Intracortical recording and manipulation of neural signals is an emerging clinical technique for diagnosing and addressing impaired brain function and treating neurological disorders^2,3^. Yet, significant unknowns in how the presence of an indwelling foreign object alters the structure and physiology of various brain cells, and by extension the robust and reliable recording of extracellular potentials, limit the long-term clinical applications of the technology^4–7^. Furthermore, current standard operating procedures do not presently screen clinical users of implantable neural interfaces for potential history of genetic predisposition to neurodegenerative or neurological disease, presenting a significant confound toward current efforts in understanding the brain’s immune response to neural electrode implants.

Implantation of intracortical microelectrodes can have long-lasting negative effects on neuronal health and brain function. Insertion of a stiff, sharp microelectrode can immediately sever neurons, axons, and blood vessels within the parenchyma, generating mechanical strain, neuroinflammation, and cerebral bleeding^5,8,9^. This acute insult leads to the activation of local glial and immune cells and release of factors which promotes inflammation, excitotoxicity, and oxidative stress, compromising local metabolic support^4^. The long-term presence of an indwelling electrode inevitably results in the development of an encapsulating glial scar impermeable to the diffusion of ions and metabolites required to detect neuronal activity, which can physically displace neurons farther from the device surface and produce a neurotoxic microenvironment^5,10^. The gradual decline in performance and fidelity of recording and stimulating electrodes is ultimately attributed to this seemingly unavoidable progression in gliosis and neurodegeneration^11^. However, attempts at identifying biological correlates of signal degradation, such as glial activation and neuronal viability, have proven unsuccessful, suggesting other biological mechanisms may be responsible for brain tissue response outcomes and long-term device performance^5^.

Age-related degenerative brain diseases, such as Alzheimer’s disease (AD), culminate in late-stage neuron loss and chronic tissue inflammation on account of presently unknown neurodegenerative mechanisms which gradually worsen over time^12,13^. AD is characterized by the appearance of extra-synaptic amyloid beta (Aβ) plaques and intra-neuronal neurofibrillary tau tangles and is clinically diagnosed by functional impairments in cognition and behavior. Despite several clinical trials demonstrating promising therapeutic potential at reducing Aβ and tau pathology or ameliorating clinical symptoms^13,14^, there is currently no known cure or effective treatment for AD. The inherent biological cause(s) of AD remain elusive but several genetic (e.g. *APOE, APP, TREM2*)^15–17^ and non-genetic^18^ (e.g. diet, exercise, cardiovascular health, and brain injury) risk factors for developing AD have been proposed. Particularly, trauma suffered to the brain has been linked to an increased risk of AD, years after the initial insult, and is most likely due to the nature in which brain injury and dementia share similar patterns of neurodegeneration^19^. The progression of AD as it is currently known shares many similarities with that of focal injury from a penetrating microelectrode array such as neuroinflammation, glial activation, and vascular injury. Hyperphosphorylated tau, a precursor for development of pathological tau tangles observed in AD, is reportedly elevated near chronically implanted microelectrodes within both wild-type rats and human Parkinsonian patients^20^. It is currently unknown, however, whether the neurotrauma suffered by intracortical microelectrodes in healthy control subjects is in any way similar to that observed in AD progression.

Characterizing Aβ deposition and tau accumulation in Alzheimer’s and other neurodegenerative diseases often require the use of genetically specialized and aged mouse models, a cost- and time-expensive process. Furthermore, under normal conditions it is often difficult to pinpoint the time and location of the biological processes responsible for the onset and progression of neurodegeneration. In contrast, focal tissue injury due to insertion of a penetrating microelectrode occurs at a known time and location within the brain, which allows for neurodegenerative processes to be spatiotemporally characterized from the onset. In this study, we investigated the appearance, if any, of age- and disease-associated factors following chronic implantation (12-16 weeks) in healthy, wild-type mice as well as a genetically susceptible mouse model of Alzheimer’s disease (APP/PS1). We hypothesized that the foreign body response to intracortical electrodes promotes the onset and progression of brain pathology typically observed within aging and degenerative brain disease, resulting in long-term neuron loss. Using *in vivo* imaging and post-mortem immunohistology, we determined that microelectrode implants exacerbate the accumulation of aberrant organelles and proteins such as lipofuscin and amyloid and promote the expression of other pathological factors associated with degenerative brain disease. In summary, our work improves our understanding of neurodegenerative processes at the intersection of brain injury and disease, which will have profound impacts on development of future intervention strategies for both implantable neural interfaces and neurological disorders within the brain.

## 3.0. METHODS

### 3.1. Experimental animal models

C57BL/6J (2 months old and 18 months old, male, 22-30g, strain# 664, Jackson Laboratories; Bar Harbor, ME), B6.Cg-Apoe^tm1.1(APOE*4)Adiuj^App^em1Adiuj^Trem2^em1Adiuj^/J (18 months old, male, 22-30g, strain# 30670, Jackson Laboratories; Bar Harbor, ME), and APP/PS1 (2 months old and 6 months old, male, 22-30g, strain# 34829, Jackson Laboratories; Bar Harbor, ME) were used in this study. All animal care and procedures were performed under approval of the University of Pittsburgh Institutional Animal Care and Use Committee and in accordance with regulations specified by the Division of Laboratory Animal Resources.

### 3.2. Probe implantation surgery

WT and AD mice (6 and 18 m.o.) were implanted with a four-shank Michigan style microelectrode array for awake, head-fixed imaging, as described previously^5,8,10,21–23^. Briefly, mice were sedated with an anesthetic cocktail (7 mg/kg xylazine and 75 mg/kg ketamine). The surgical site was shaved and sterilized 2-3 times with alternate scrubs of an aseptic wash and 70% ethanol. Mice were then fixed to a stereotaxic frame and a small incision was made over the skull. Care was given to remove all skin, hair, and connective tissue from the surface of the skull before a thin layer of Vetbond (3M) was applied to the surface. A rectangular head bar was fixed to the skull for awake, head-fixed imaging. Four bone screw holes were drilled and bone screws inserted over both ipsilateral and contralateral motor cortices and lateral visual cortices to secure the head bar. A 4 mm by 4 mm bilateral craniotomy was performed prior to probe insertion. The skull was periodically bathed in sterile saline to prevent the skull from overheating during drilling. Probes were inserted through an intact dura into the cortex at a 30° angle at 200 μm/s for a total linear distance of ~600 μm (oil hydraulic Microdrive; MO-82, Narishige, Japan) and final z-depth of ~250-300 μm beneath the pial surface. The craniotomy was filled with sealant (Kwik-Sil) before sealing with a glass coverslip and dental cement. Anesthesia was maintained throughout the surgery with ketamine (40 mg/kg), as needed. For immunohistochemistry, a separate cohort of C57BL/6J and APP/PS1 mice were implanted as described above with the exception of a single-shank probe implanted at a 90° angle, as described previously^7^. Briefly, a drill-bit sized craniotomy was formed over the left primary visual (V1) cortex (1.5 mm anterior and 1 mm lateral from lambda) and a Michigan-style microelectrode was inserted at a rate of 200 μm/s to a final resting depth of 1600 μm below the pial surface. The probe and craniotomy were filled with Kwik-sil and a headcap was secured with dental cement. Ketofen (5 mg/kg) was provided post-operatively up to two days post-surgery or as needed.

### 3.3. Two-photon imaging and Aβ labeling

Two-photon microscopy was used to track the rate of amyloid deposition or clearance around implanted microelectrodes in 6 month old WT and APP mice over a 16-week implantation period (0, 2, 4, 7, 14, 21, 28, 56, 84, and 112 days post-implantation), as described previously^8,10,21,22^. The microscope was equipped with a scan head (Bruker, Madison, WI), a OPO laser (Insight DS+; Spectra-Physics, Menlo Park, CA), non-descanned photomultiplier tubes (Hamamatsu Photonics KK, Hamamatsu, Shizuoka, Japan), and a 16X, 0.8 numerical aperture water-immersive objective lens (Nikon Instruments, Melville, NY). To visualize amyloid, mice were injected intraperitoneally with methoxy-X04 (2 mg/kg, Abcam, #ab142818) 24 hours prior to imaging^20^. Mice were retro-orbitally injected with FITC-dextran (2 MDa, 0.03 mL at 10 mg/mL) immediately prior to imaging to visualize surrounding blood vessels. The vasculature was used as a landmark to ensure similar ROI fields were captured between subsequent imaging sessions. Methoxy-X04 was excited at a 740 nm laser excitation wavelength and care was given not to exceed >20-30 mW of power during chronic imaging. Z-stacks were collected along the full depth of the implant at a step size of 2-3 μm, ~5 s frame rate, and zoom factor of ~1.5-2x.

### 3.4. Immunohistochemical staining

C57BL/6J and APP/PS1 mice that were implanted at ages of 2 months old were sacrificed and perfused according to University of Pittsburgh IACUC approved methods at 1 week (*n* = 6 per group) or 16 weeks (*n* = 7 per group) post-implantation, as described previously^7^. Briefly, mice were sedated using a cocktail mixture of xylazine (7 mg/kg) and ketamine (75 mg/kg). A toe-pinch test was performed to ensure proper level of anesthesia prior to beginning the procedure. For each mouse, 100 mL of warm phosphate buffered saline (PBS) was perfused transcardially (pump pressure between 80-100 mm Hg) followed by 100 mL of 4% paraformaldehyde (PFA). Mice were then decapitated with both skull and implant left intact for 24 h post-fixation at 4°C in 4% PFA. Brains were then carefully detached from both the skull and implant and sequentially soaked in 15% and 30% sucrose in PBS at 4°C for 24 h each. Once the brain samples had reached sucrose equilibration, they were frozen in a 2:1 ratio of 20% sucrose:optimal cutting temperature compound (Tissue Tek, Miles Inc., Elkhart, IN, United States). Frozen brain samples were then sectioned horizontally using a cryostat (Leica Biosystems, Wetzlar, Germany) at a 25 μm thickness throughout the entire depth of the implant (~1600 μm).

Before staining, frozen tissue sections were re-hydrated with two washes of 1x PBS for 5 min each. For antigen retrieval, slides were incubated in 0.01 M sodium citrate buffer for 30 min at 60°C. Slides were then incubated in in a peroxidase blocking solution (10% v/v methanol and 3% v/v hydrogen peroxide in 1x PBS) for 20 min on a table shaker (60 r.p.m.) at room temperature (RT) to block for active aldehydes and reduce the chance of non-specific binding. Sections were then pre-treated with a solution of 1% triton X-100 and 10% donkey serum in 1x PBS for 1 hr at RT followed by blocking endogenous mouse immunoglobulin G (IgG) with donkey anti-mouse IgG fragment (Fab) for 2 h at 1:10 dilution at RT. Then, sections were rinsed with alternating washes of 1x PBS and 1x PBS-T (1% v/v of Tween-20 in 1x PBS) for 4 times 4 min each. Primary antibodies were diluted in solution of 1% triton X-100 and 10% donkey serum and applied to slides for 12-18 hr at 4°C. All primary antibodies used in this study are listed in Table 1. Sections were rinsed in 3×5 min washes of 1x PBS the following day.

**Table 1.** List of primary antibodies used for immunohistochemical staining.

| Primary antibody | Target | Supplier | Host | Dilution |
| --- | --- | --- | --- | --- |
| Anti-6E10 | Amyloid beta | Enzo Life Sciences (ABS612) | Rb | 1:200 |
| Anti-APP | Amyloid precursor protein | Abcam (ab2084) | Gt | 1:400 |
| Anti-AT8 | Phospho-tau | Thermo Fisher (MN1020B) | Biotin | 1:250 |
| Anti-GFAP | Astrocytes | Abcam (ab4674) | Ck | 1:500 |
| Anti-Iba1 | Microglia | Sigma (MABN92) | Ms | 1:400 |
| Anti-Myelin Basic Protein | Myelin | Abcam (ab7349) | Rt | 1:100 |
| Anti-NeuN | Neurons | Thermo Fisher (MA5-33103) | Ms | 1:500 |
| Anti-NF200 | Neurofilament | Sigma (N5389) | Ms | 1:250 |
| Anti-Trem2 | Triggering receptor expressed on myeloid cells 2 | R&D Systems (AF1729) | Sh | 1:100 |

Secondary antibodies were diluted 1:500 in 1x PBS and applied to slides for 2 hr at RT. Secondary antibodies used were: Alexa Fluor 488 donkey anti-mouse, Alexa Fluor 488 donkey anti-rabbit, Alexa Fluor 568 donkey anti-mouse, Alexa Fluor 568 donkey anti-rat, Alexa Fluor 568 donkey anti-goat, Alexa Fluor 568 donkey anti-sheep, Alexa Fluor 647 donkey anti-mouse, Alexa Fluor 647 donkey anti-chicken, and Alexa Fluor 647 streptavidin (Abcam, Cambridge, UK). Then, slides were rinsed once more with 3×5 min washes of 1x PBS. To stain for cell nuclei, slides were incubated in Hoechst 33342 (Invitrogen) at 1:1000 in 1x PBS for 10 min at RT followed by another rinse 3×5 min with 1x PBS. Finally, slides were cover slipped using Fluoromount-G (Southern Biotech, Birmingham, AL, United States) and sealed prior to imaging.

Samples were imaged using a confocal microscope (FluoView 1000, Olympus, Inc. Tokyo, Japan) using a 20x oil-immersive objective lens (Nikon Instruments, Melville, NY). For each section, an ipsilateral and contralateral image was captured using same laser intensity and image settings for data normalization. For each image, a z-stack was collected consisting of 6 image planes each spaced 5 μm apart (635.9 x 635.9 μm, 1024 x 1024 pixels on FV10-ASW Viewer V4.2). Raw images were saved as 16-bit grayscale TIFFs.

### 3.5. Data analysis

For two-photon quantification, the distance between lipofuscin granules or amyloid plaques and the implant were determined by referencing to a 3D distance map of the electrode surface, as described previously^5^. First, a 2D mask of the electrode footprint was manually outlined within ImageJ (National Institute of Health). Considering the original electrode implantation angle of 25-30° this 2D electrode mask was digitally rotated using ImageJ’s built-in ‘Interactive Stack Rotation’ tool. A distance map was then generated by applying a distance transform to this 3D mask using the ‘bwdist’ function in Matlab (MathWorks, Boston, MA). The spatial coordinates of segmented lipofuscin granules or amyloid plaques were cross-referenced with this 3D distance map to determine the nearest Euclidean distance to the electrode surface for each timepoint.

For lipofuscin quantification, lipofuscin granules were identified by their natural autofluorescence signal detected across multiple emission filters (595nm for green and 525nm for red, 50nm bandpass). First, multi-channel z-stack images were spectrally unmixed using the ImageJ plugin, ‘LUMoS Spectral Unmixing’. Then, background noise was subtracted by applying a Gaussian filter (sigma = 20) to the unmixed image containing the lipofuscin signal and subtracting the filtered image from the original. Background was further reduced by applying a bandpass filter, followed by a median filter, then finally using ImageJ’s ‘Despeckle’ function. The resultant image was binarized by applying a fluorescence intensity threshold (mean + 2.5 standard deviation) before running the ‘3D Objects Counter’ plugin to determine the size and spatial coordinates of segmented lipofuscin granules. For amyloid quantification, images with methoxy-X04 signal and 6E10 signal were processed similarly as described above. The spatial coordinates of both amyloid clusters and plaques were referenced to respective 3D electrode distance map for each timepoint. The rate of change in size of individual amyloid plaques were tracked longitudinally by referencing with the vasculature as a landmark.

For fluorescence intensity analyses of stained tissue, images were processed using a semi-automated intensity-based custom Matlab script (INTENSITY Analyzer), as described previously^7,24^. Briefly, images were binned concentrically every 10 μm up to 300 μm away from the probe center. The average grayscale intensity was calculated for every pixel above a threshold determined from the intensity of the background noise for each bin. Fluorescence intensities were then normalized using average intensity values collected from the contralateral hemisphere of the section (i.e., intact hemisphere). For cell counting of stained tissue, images were binned concentrically every 50 μm up to 300 μm away from the probe center. Cells were identified and counted based on the presence of DAPI-stained nuclei and data was similarly normalized using cell counts from the contralateral hemisphere. Normalized intensity and cell counts were average over all animals per timepoint per group and presented as mean ± standard error.

### 3.6. Statistics

A two-way repeated measures ANOVA (*p* < 0.05) was used to determine significant differences in changes in plaque volume, staining fluorescence intensities, and histological cell counts between WT and APP/PS1 mice followed by a post-hoc t-test with a Bonferroni correction to determine pairwise significances between groups. A Welch’s t-test (*p*<0.05) was used to determine significant differences in 6E10 staining plaque area.

## 4.0. RESULTS

Unlike humans, rodents do not develop Alzheimer’s pathology naturally with aging. Therefore, specially developed transgenic mouse models are required to study the progression of neuropathology within Alzheimer’s disease^25^. Here, we use the APP/PS1 mouse model expressing a humanized form of the *APP* gene, which promotes overexpression of the amyloid precursor protein, as well as the *PSEN1* gene, which controls for expression of the enzyme presenilin 1 required for proteolytic cleavage of amyloid (**Fig. 1a**). These mice have been extensively characterized for study of Alzheimer’s disease and therefore their temporal onset and progression of amyloid pathology is well known^26^. To determine whether chronic electrode implantation leads to accumulation of age- or disease-associated factors, we used two-photon microscopy to longitudinally quantify aberrant protein aggregation around a multi-shank Michigan style microelectrode array within the WT and AD mouse cortex (**Fig. 1b**). Specifically, to assess age-related lipofuscin accumulation, mice were aged to 18 months prior to electrode implantation surgery. Next, to assess for amyloid deposition, young (2-month-old) and adult (6-month-old) APP/PS1 mice were intraperitoneally administered methoxy-X04 (MX04), a Congo-red derivative which binds to amyloid and tau, 24 hrs prior to each imaging session (**Fig. 1c**).

**Figure 1.**
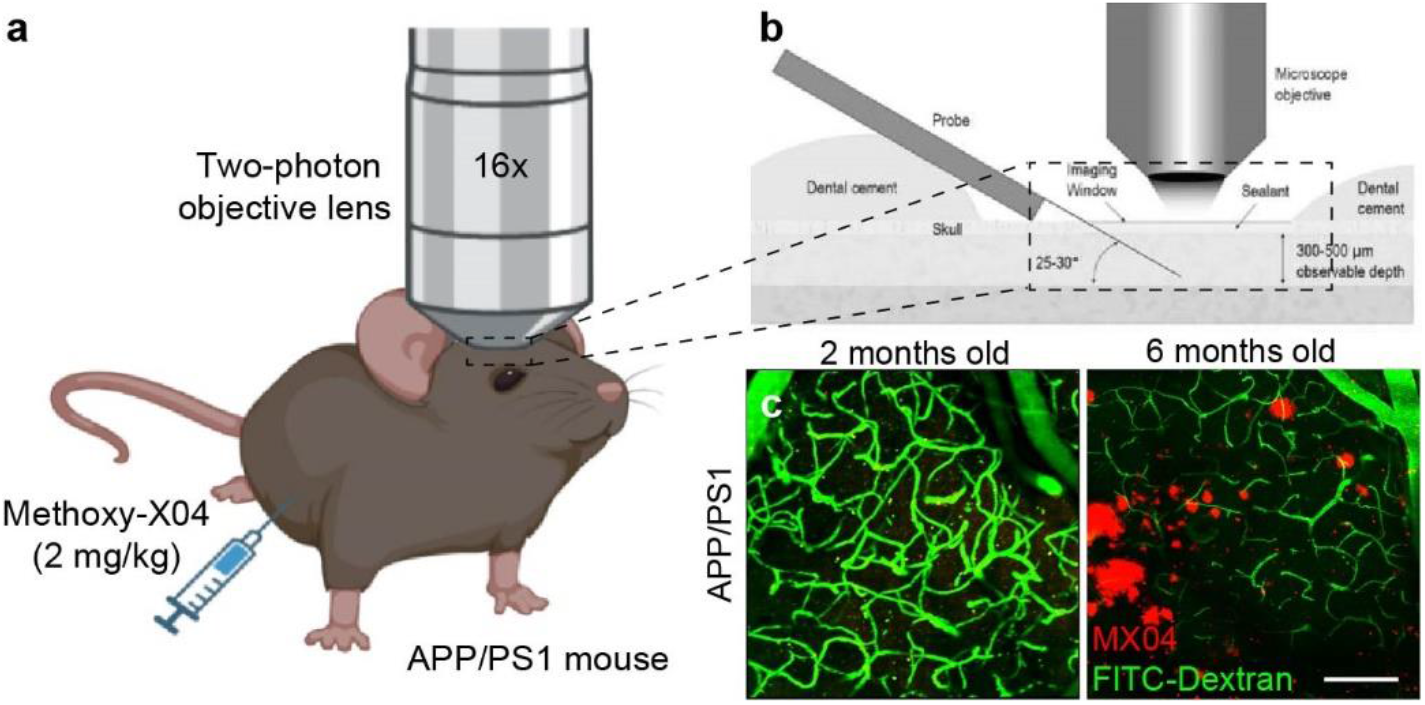
Two-photon visualization of age- and disease-related factors in a mouse model of Alzheimer’s disease. (a) Methoxy-X04 (2 mg/kg) was administered to APP/PS1 mice in order to visualize amyloid deposition within the mouse brain using two-photon microscopy. (b) Schematic of microelectrode implantation within the mouse cortex underneath an optical window for longitudinal imaging of chronic tissue response. (c) Representative two-photon image demonstrating age-dependency of plaque visualization (MX04, *red*) in APP/PS1 mice. Vasculature is labeled with FITC-dextran (*green*). Scale bar = 100 μm.

### 4.1. Aberrant accumulation of lipofuscin around chronically implanted electrodes in aged WT and AD mice

Lipofuscin is an aging pigment with natural auto-fluorescent properties that can be imaged without the need for transgenic models or additional fluorophore labeling using 1:1 fluorescent signal detected across multiple wavelength emission filters (green and red channels) using two-photon microscopy (**Fig. 2a**). These lipofuscin form when lysosomes are unable to fully degrade and recycle cellular waste due to insufficient pH and lysosomal hydrolysis^27–29^. To assess whether chronic microelectrode implantation leads to aberrant protein aggregation with aging or in neurodegenerative disease, lipofuscin was assessed over a 12-week implantation period in an 18-month-old AD mouse model (hAbeta/APOE4/Trem2*R47H). The hAbeta/APOE4/Trem2*R47H mouse is a triple mutant strain carrying a humanized *ApoE* knock-in mutation, a humanized *App* allele, and a *Trem2* mutation. Accumulation of lipofuscin was readily observed over time around an implanted microelectrode array in both the hAbeta/APOE4/Trem2*R47H model and an age-matched wild-type mouse (**Fig. 2b**). Lipofuscin granules increased in volume and accumulated in larger densities with closer proximity to the implant surface (**Fig. 2c-f**). These results demonstrate that chronic microelectrode arrays can exacerbate aberrant protein aggregation near the site of implantation.

**Figure 2.**
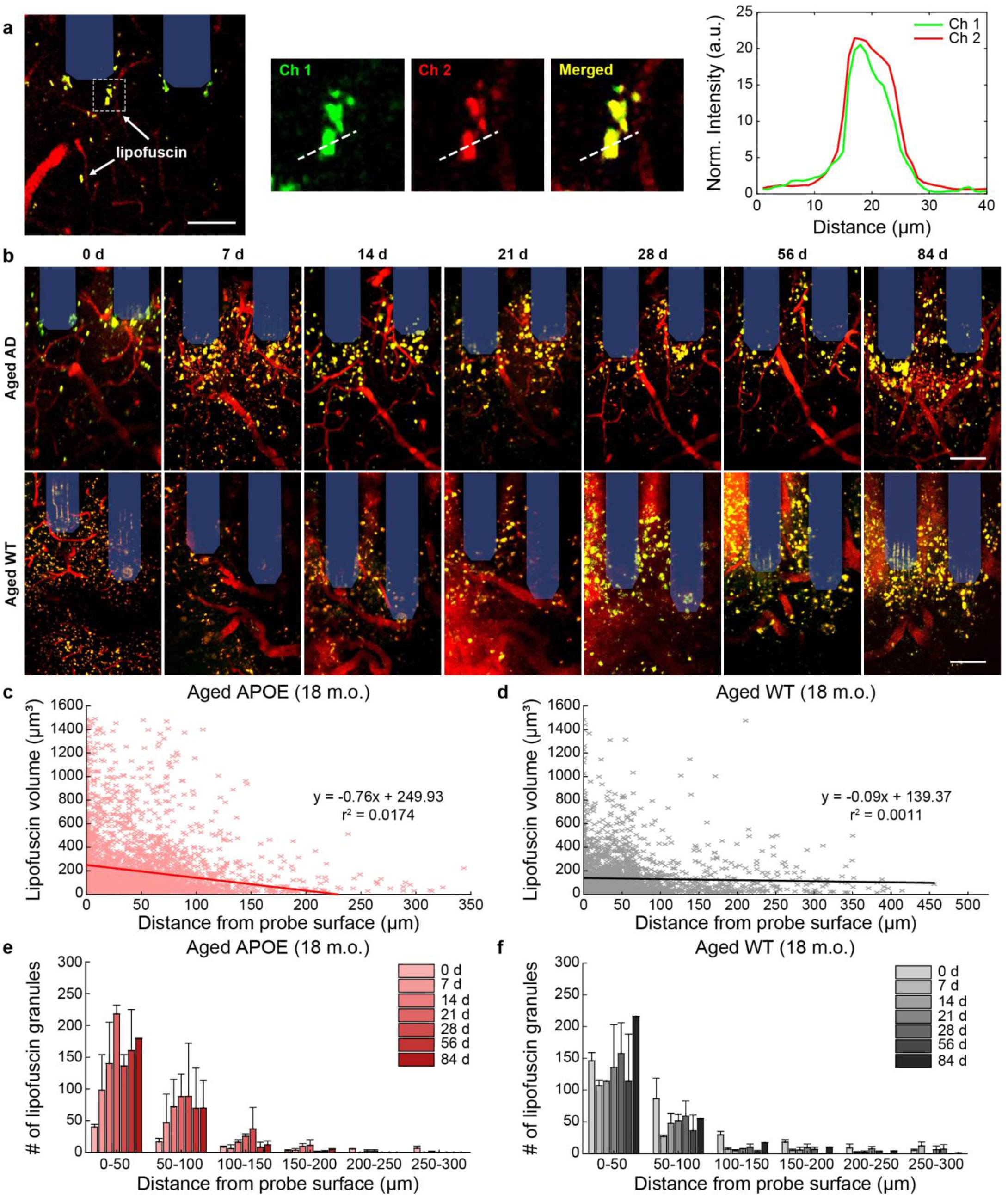
Lipofuscin accumulation around chronically implanted microelectrodes in aged WT and AD mice. (a) Example two-photon image of lipofuscin signal detected across multiple wavelength emission filters. Lipofuscin can be detected by a 1:1 fluorescence intensity overlap in both red and green filtered channels. (b) Representative two-photon images of accumulated lipofuscin granules (*yellow*) around multi-shank microelectrode array (*shaded blue*) over 12-week post-implantation in aged (18 m.o.) WT and AD mice. Scale bars = 50 μm. (c-d) Scatter plots demonstrating trend in volume of lipofuscin granules with respect to distance from probe surface (aged APOE: 3,173 lipofuscin granules across 7 timepoints; aged WT: 2,757 lipofuscin granules across 7 timepoints). (e-f) Average number of lipofuscin granules with binned distance from the probe in aged AD and WT mice (*n* = 2 mice per group). All data is reported as mean ± SEM.

### 4.2. Chronic electrode implantation halts the growth of pre-existing Aβ plaques while promoting local accumulation of new amyloid clusters

Amyloid beta (Aβ) is another protein whose aggregation and insufficient clearance from the brain is implicated in aging and neurodegenerative disease. To understand whether device implantation injury impacts the morphology of pre-existing Aβ plaques within the brain, we inserted microelectrodes within 6-month-old APP/PS1 mice and longitudinally assessed the growth of nearby Aβ plaques using intravenously administered MX04 over a 12-week implantation period. At this age, APP/PS1 mice readily present Aβ plaques throughout the cortex and therefore provide the opportunity assess changes in volume of pre-existing plaques around chronically implanted microelectrodes (**Fig. 3a**). In the uninjured cortex of APP/PS1 mice, Aβ plaques begin to manifest around 5 months of age and continually increase in size until around 10-12 months^22^. Surprisingly, we reveal that the Aβ plaques located on the ipsilateral hemisphere around chronically implanted microelectrodes do not change in size with chronic implantation compared to plaques on the contralateral hemisphere, whose volumes were significantly increased (**Fig. 3b**, *p*<0.05, two-way ANOVA). We also show that the plaques quantified on the ipsilateral hemisphere near implanted microelectrodes decrease in size over a 12-week implantation period whereas plaques distal from the electrode demonstrate a significant increase in percent change in volume during the implantation period (**Fig. 3c**, *p*<0.05, two-way ANOVA).

**Figure 3.**
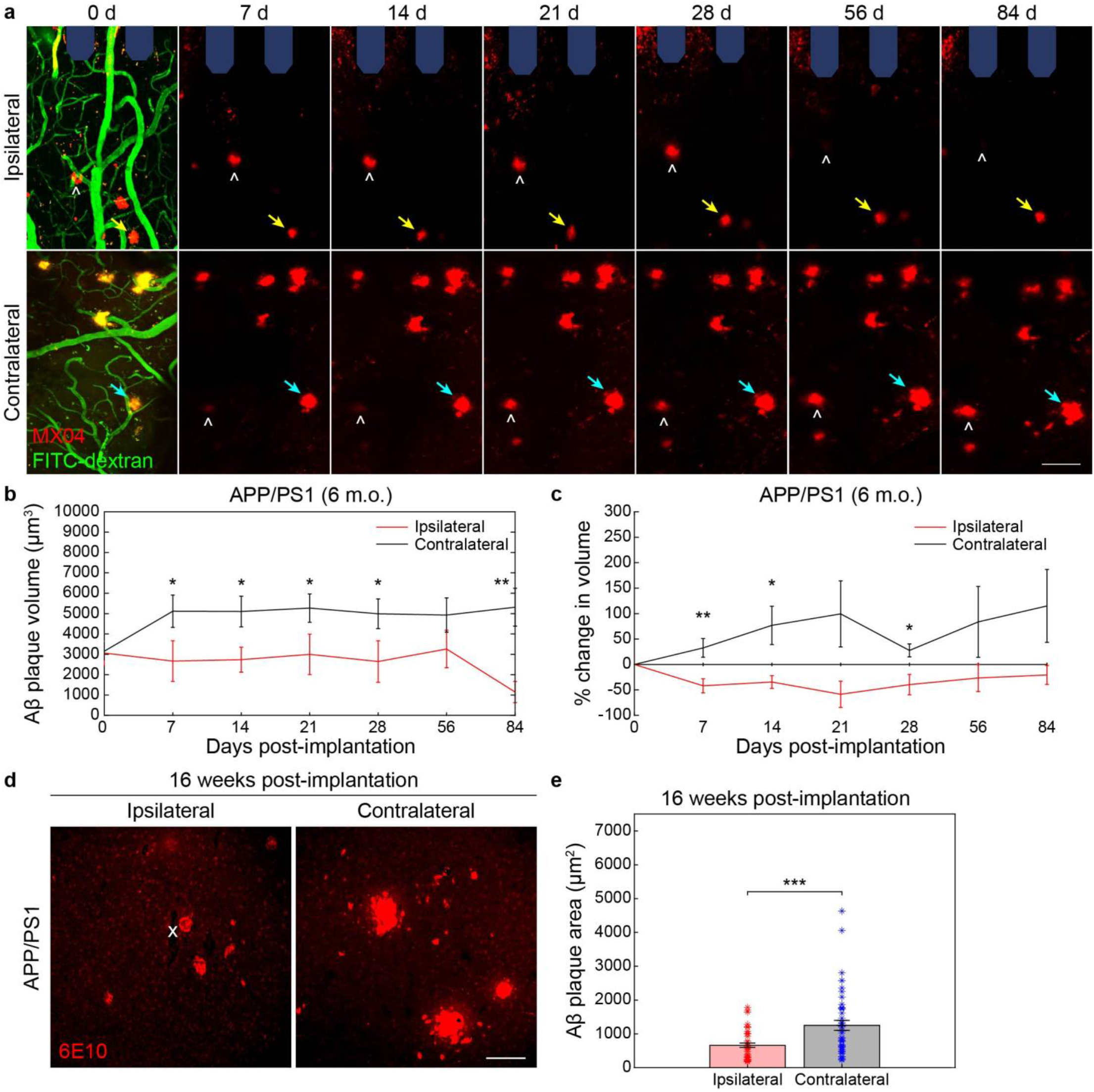
Chronic microelectrode implantation reduces the growth of local amyloid plaques in adult APP/PS1 mice. (a) Representative two-photon images of Aβ plaques labeled with methoxy-X04 (MX04, *red*) and blood vessels (FITC-dextran, *green*) in ipsilateral hemisphere around multi-shank microelectrode array (*shaded blue*) over 12 weeks post-implantation in adult (6 m.o.) APP/PS1 mice compared to contralateral (uninjured) hemisphere. Ipsilateral hemisphere demonstrates Aβ plaques which do not visually change in size with chronic implantation (*yellow arrow*) whereas Aβ plaques on the contralateral hemisphere appear to increase in size over time (*cyan arrow*). NOTE: some plaques move into and out of frame over time (*white hat*) due to tissue drift between subsequent chronic imaging sessions. Scale bar = 50 μm. (b) Change in Aβ plaque volume over a 12 week implantation period between ipsilateral and contralateral hemispheres in adult APP/PS1 mice (25 Aβ plaques on ipsilateral hemisphere and 27 Aβ plaques on contralateral hemisphere tracked longitudinally over 7 time points across *n* = 3 mice). (c) Percent change in Aβ plaque volume with respect to plaque size on day 0 of electrode insertion over a 12-week implantation period between ipsilateral and contralateral hemispheres in adult APP/PS1 mice. (d) Representative immunohistology stain for 6E10, an Aβ marker, following 16 weeks post-implantation in adult (6 m.o.) APP/PS1 mice demonstrating visually reduced Aβ plaque sizes in ipsilateral hemisphere around the site of probe insertion (denoted by white ‘x’) compared to contralateral side. Scale bar = 100 μm. (e) Average Aβ plaque area measured by 6E10 stain between ipsilateral and contralateral hemispheres (*n* = 42 Aβ plaques on ipsilateral hemisphere over 13 histological tissue sections and *n* = 45 Aβ plaques on contralateral hemisphere over 17 histological sections across 6 mice total). * *p* < 0.05, ** *p* < 0.01, *** *p* < 0.001. All data is reported as mean ± SEM.

In support of our *in vivo* imaging results, we demonstrate via immunohistochemical staining of post-mortem tissue for 6E10, a common AD marker used to label amyloid beta, that amyloid plaques on the ipsilateral hemisphere are considerably smaller than plaques found on the contralateral hemisphere following 16-weeks post-implantation in APP/PS1 mice (**Fig. 3d**). Quantifying the stained 6E10 area of individual Aβ plaques, we determine that plaques on the ipsilateral hemisphere are, on average, significantly smaller than those located on the contralateral hemisphere further away (**Fig. 3e**, *p*<0.001, Welch’s t-test). Altogether, these findings suggest that device implantation injury effectively inhibits the growth of Aβ plaques located around chronically implanted microelectrodes.

To determine whether microelectrode implantation preferentially favors the accumulation of amyloid, we first inserted microelectrode arrays within 2-month-old APP/PS1 mice. APP/PS1 mice at this age have yet to present stereotypical amyloid pathology within the brain and therefore provide an opportunity to assess whether device injury triggers the aggregation of amyloid protein prior to the temporally defined onset of neuropathology within the AD model. It is also important to note that these mice do not yet present lipofuscin pigments within the brain at this age as well. Surprisingly, we observed the appearance of MX04-labeled amyloid deposits, in which we term here as “amyloid clusters”, within just the first 7 days following microelectrode implantation in young APP/PS1 mice (**Fig. 4a**). From our analyses, we reveal a slight negative association between the size of Aβ clusters and their individual distances from the surface of an implanted microelectrode array (**Fig. 4b**, R^2^ = 0.0123). Additionally, we determined that the number of quantified Aβ clusters increases preferentially with distance near the implant and over time following chronic microelectrode implantation (**Fig. 4c**). These results suggest that the tissue injury sustained from chronic electrode insertion within the brain accelerates the accumulation of amyloid pathology weeks to months before the expected onset of neuropathology within an uninjured mouse model of Alzheimer’s disease.

**Figure 4.**
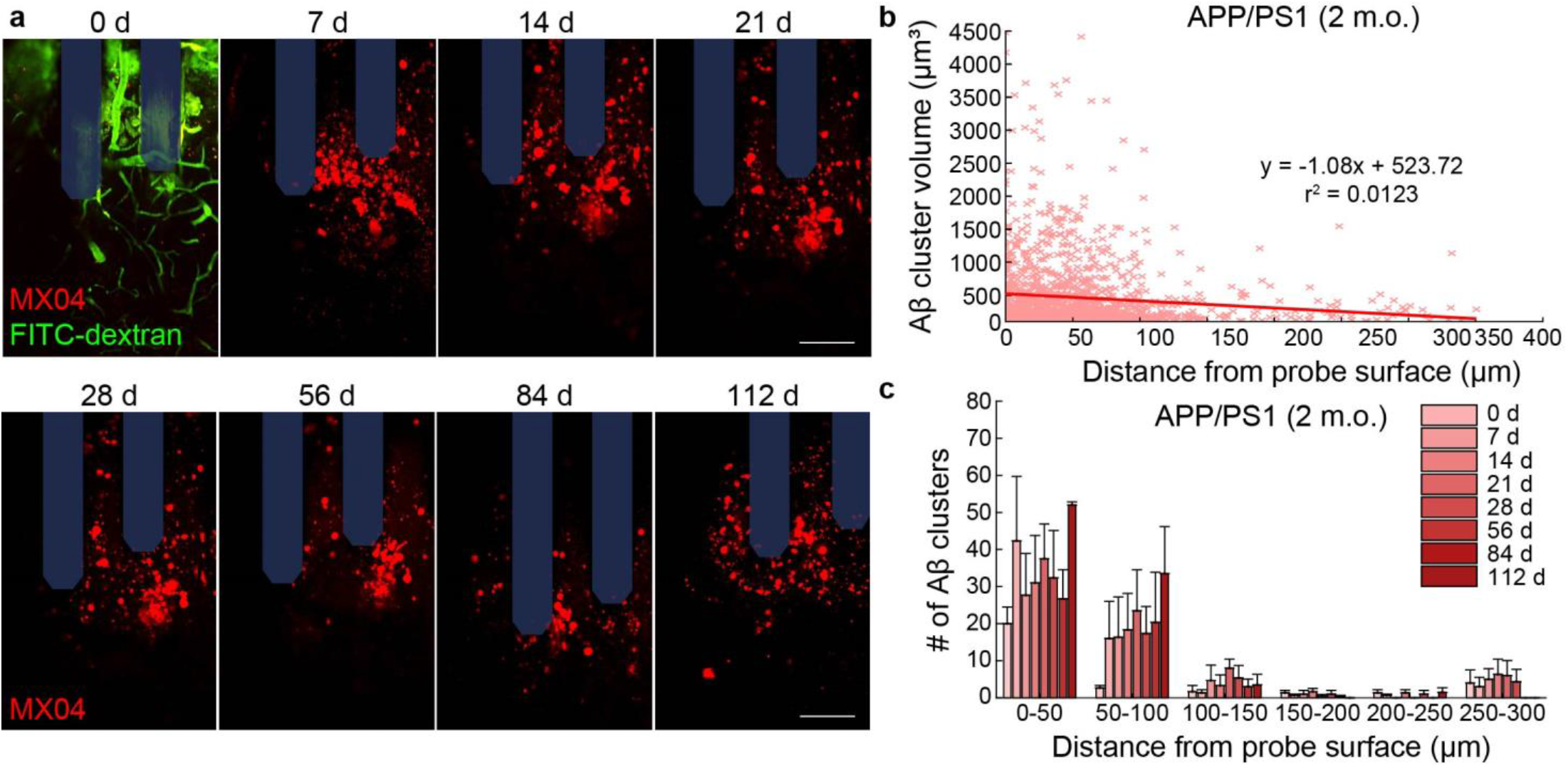
Accumulation of amyloid clusters around chronically implanted microelectrodes in young APP/PS1 mice. (a) Representative two-photon images of Aβ clusters labeled with methoxy-X04 (MX04, *red*) and blood vessels (FITC-dextran, *green*) around multi-shank microelectrode array (*shaded blue*) over 16 weeks post-implantation in young (2 m.o.) APP/PS1 mice. Scale bars = 50 μm. (b) Scatter plot demonstrating a trend in increased volume of Aβ clusters with respect to distance from probe surface (1,407 Aβ clusters across 7 timepoints). (c) Average number of Aβ clusters with binned distance from the probe in young APP/PS1 mice (*n* = 3). All data is reported as mean ± SEM.

### 4.3. Elevated phagocytosis in activated microglia and amyloid precursor protein in reactive astrocytes around chronically implanted electrodes

Microglia assume a critical role in phagocytosing cellular and tissue debris following brain injury or amyloid protein in Alzheimer’s disease and their dysfunction has been linked to the progression of neurodegenerative disease^30,31^. To determine whether phagocytosis within microglia due to device injury is elevated around implanted microelectrodes or within AD, explanted WT and APP/PS1 brain tissue were stained for Iba-1, a microglial marker, and ‘triggering receptor expressed on myeloid cells 2’ (TREM2), an innate immune receptor predominantly expressed by microglia within the brain during conditions of metabolic stress^25^. Interestingly, we found that microglia near the site of electrode implantation strongly co-localize with the expression of TREM2 (**Fig. 5a**). As expected, Iba-1 fluorescence intensities were visually elevated near the site of probe implantation at 1- and 16-weeks post-implantation in WT and APP/PS1 mice (**Fig. 5b**). While normalized Iba-1 fluorescence intensities were increased with proximity to the site of electrode implantation, we did not reveal any significant differences in Iba-1 fluorescence intensities between WT and APP/PS1 at either 1-week or 16-week post-implantation (**Fig. 5c,d**). Interestingly, we also observed a visual increase in TREM2 fluorescence intensities near the site of probe implantation at 1- and 16-weeks post-implantation in WT and APP/PS1 mice (**Fig. 5e**). While normalized TREM2 fluorescence intensities were increased closer to the site of implantation, we did not reveal any significant differences in TREM2 fluorescence intensities between WT and APP/PS1 at either 1-week or 16-week post-implantation (**Fig. 5f,g**). Nevertheless, these findings demonstrate that activated microglia increase expression of cellular immune receptors reportedly involved in phagocytosis of cellular debris and Aβ around chronically implanted microelectrodes.

**Figure 5.**
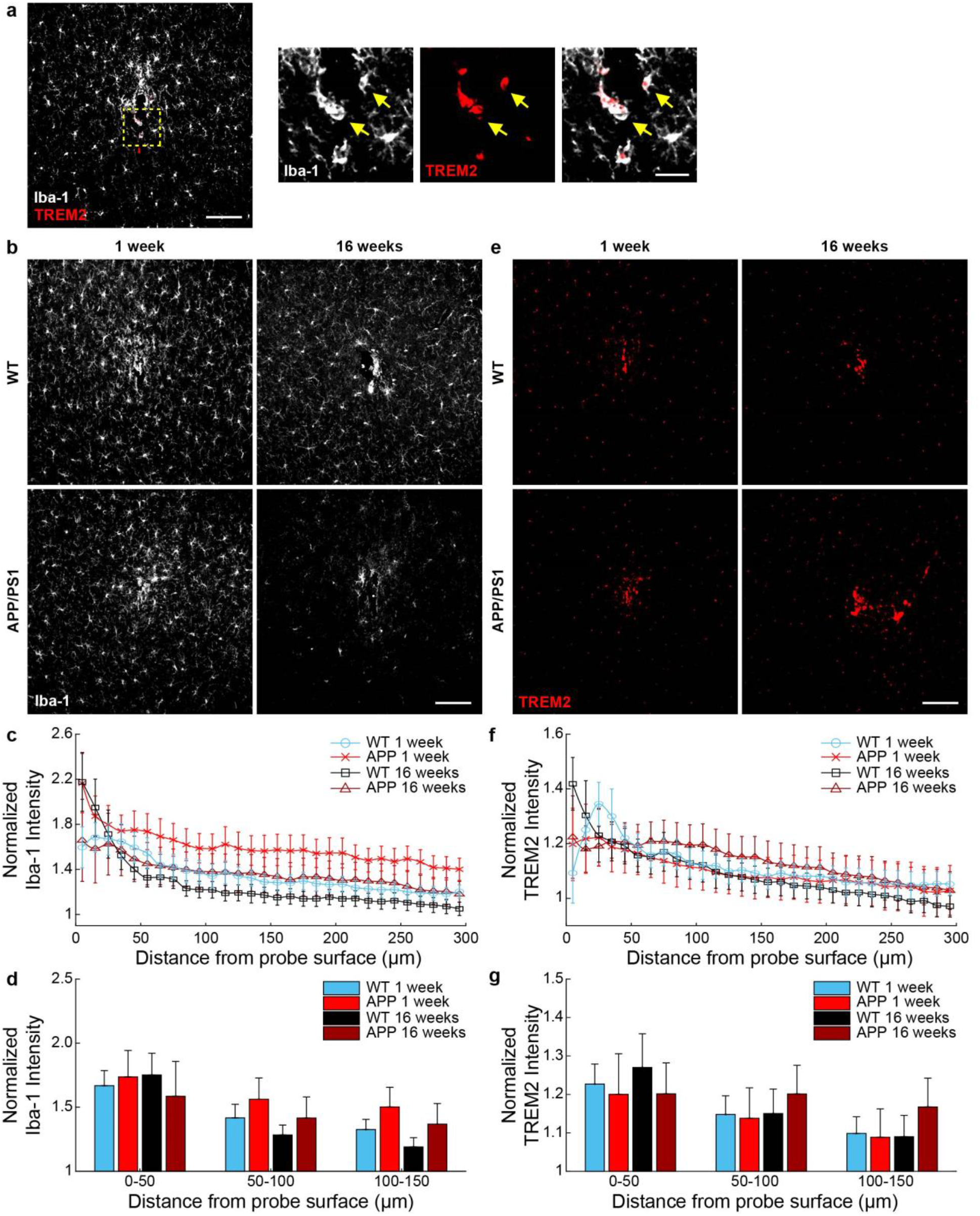
Microglial expression of phagocytic receptors around chronically implanted microelectrodes. (a) Immunohistological representation demonstrating co-localization of Iba-1+ microglia (*white*) with triggering receptor expressed on myeloid cells 2 (TREM2, *red*) around the site of electrode implantation (*yellow arrows*). Scale bar = 100 μm, 10 μm (inset). (b) Representative images of Iba-1 fluorescence staining around implanted microelectrodes at 1- and 16-weeks post-implantation in WT and APP/PS1 mice. Scale bar = 100 μm. (c) Normalized Iba-1 fluorescence intensity with respect to distance around chronically implanted microelectrodes at 1- and 16-weeks post-implantation in WT and APP/PS1 mice. (d) Average Iba-1 fluorescence intensity within 50 μm bins up to 150 μm around chronically implanted microelectrodes at 1- and 16-weeks post-implantation in WT and APP/PS1 mice. (e) Representative images of TREM2 fluorescence staining around implanted microelectrodes at 1- and 16-weeks post-implantation in WT and APP/PS1 mice. Scale bar = 100 μm. (f) Normalized TREM2 fluorescence intensity with respect to distance around chronically implanted microelectrodes at 1- and 16-weeks post-implantation in WT and APP/PS1 mice. (g) Average TREM2 fluorescence intensity within 50 μm bins up to 150 μm around chronically implanted microelectrodes at 1- and 16-weeks post-implantation in WT and APP/PS1 mice (*n* = 6 mice per group at 1 week, *n* = 7 mice per group at 16 weeks). All data is reported as mean ± SEM.

Reactive astrocytes are the main culprits in formation of an astrocytic scar around intracortical electrodes and their dysfunction is commonly reported in neurological disorders^18,26^. To assess whether astrocyte reactivity due to chronic microelectrode implantation is exacerbated in a mouse model of AD, explanted WT and APP/PS1 brain tissue were stained for GFAP, a marker for reactive astrocytes. As expected, GFAP+ astrocyte staining was markedly increased near the site of device implantation in both WT and APP/PS1 mice at 1- and 16-weeks post-insertion (**Fig. 6a**). GFAP fluorescence intensities were significantly elevated at 16-weeks post-implantation in both WT and APP/PS1 compared to 1-week post-implantation up to 150 μm from the site of electrode implantation (**Fig. 6b,c**, *p*<0.001, Welch’s t-test). Astrocytes reportedly can contribute to Aβ deposition due to their own production of amyloid precursor protein (APP)^27,28^. APP is a transmembrane protein whose pathological cleavage into aggregate forms of Aβ lead to the formation of senile plaques within AD and other dementia^29^. Interestingly, we report that GFAP+ astrocytes notably express APP near the site of probe implantation (**Fig. 6d**). This co-expression of APP was specific to astrocytes and not microglia, since APP only co-labeled with GFAP+ astrocyte cells but not with Iba-1+ microglia cells. These results suggest that device implantation injury can induce the expression of potential precursors for Aβ production within reactive astrocytes around chronically implanted microelectrodes.

**Figure 6.**
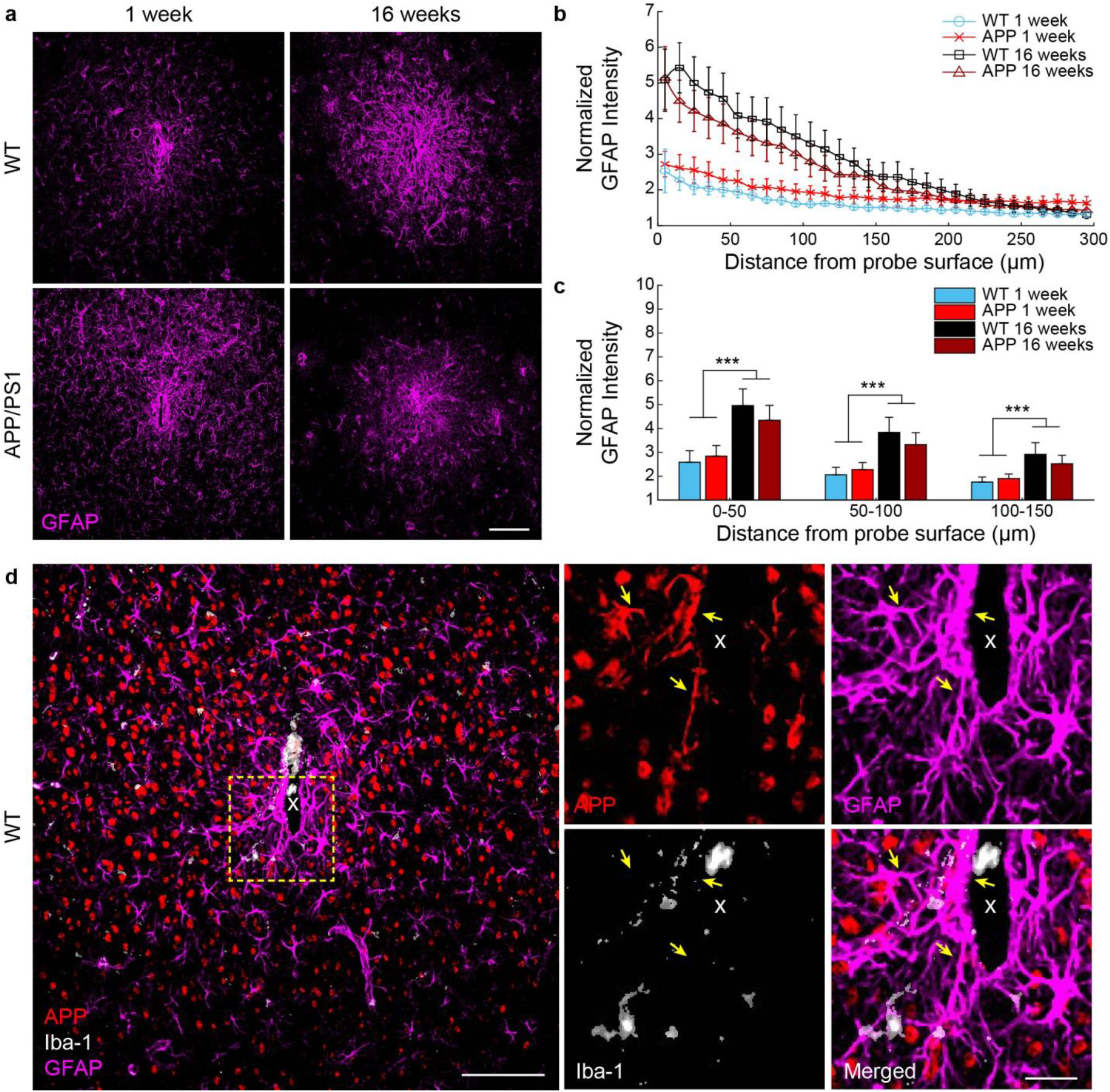
Reactive astrocytes express amyloid precursor protein around chronically implanted microelectrode arrays. (a) Representative images of GFAP+ reactive astrocyte staining (*magenta*) around implanted microelectrodes at 1- and 16-weeks post-implantation in WT and APP/PS1 mice. Scale bar = 100 μm. (b) Normalized GFAP fluorescence intensity with respect to distance around chronically implanted microelectrodes at 1- and 16-weeks post-implantation in WT and APP/PS1 mice. (c) Average GFAP fluorescence intensity within 50 μm bins up to 150 μm around chronically implanted microelectrodes at 1 - and 16-weeks post-implantation in WT and APP/PS1 mice (*n* = 6 mice per group at 1 week, *n* = 7 mice per group at 16 weeks). (d) Immunohistological example demonstrating expression of amyloid precursor protein (APP, *red*) in GFAP+ astrocytes (*magenta*) but not Iba-1+ microglia (*white*) near the site of a chronically implanted microelectrode in a WT mouse. Scale bar = 100 μm, 25 μm (inset). *** *p* < 0.001. All data is reported as mean ± SEM.

### 4.4. Neuronal densities are reduced near chronically implanted microelectrodes

We previously reported that neuronal densities are affected around chronically implanted microelectrodes up to 4 weeks post-implantation^7^. Here, we stain for NeuN to visualize the distribution of neurons around microelectrode arrays in WT and APP/PS1 mice at 1- and 16-weeks post-implantation (**Fig. 7a**). Quantification of neuronal densities normalized to contralateral hemispheres revealed a decrease in neurons within the 0-50 μm region near the site of implantation in both WT and APP/PS1 mice (**Fig. 7b**). Specifically, we determined that neuronal densities were significantly reduced in tissue regions between 0-50 μm and 100-150 μm at 1-week post-implantation and between 0-50 μm and 50-100 μm at 16-weeks post-implantation in WT mice but not APP/PS1 mice (*p*<0.05, two-way ANOVA). However, we did not detect any significant differences in neuronal density between WT and APP/PS1 mice at either 1- or 16-week post-implantation (*p>*0.05, two-way ANOVA). Nevertheless, these results reveal that chronic microelectrodes result in the loss of neurons local to the site of implantation.

**Figure 7.**
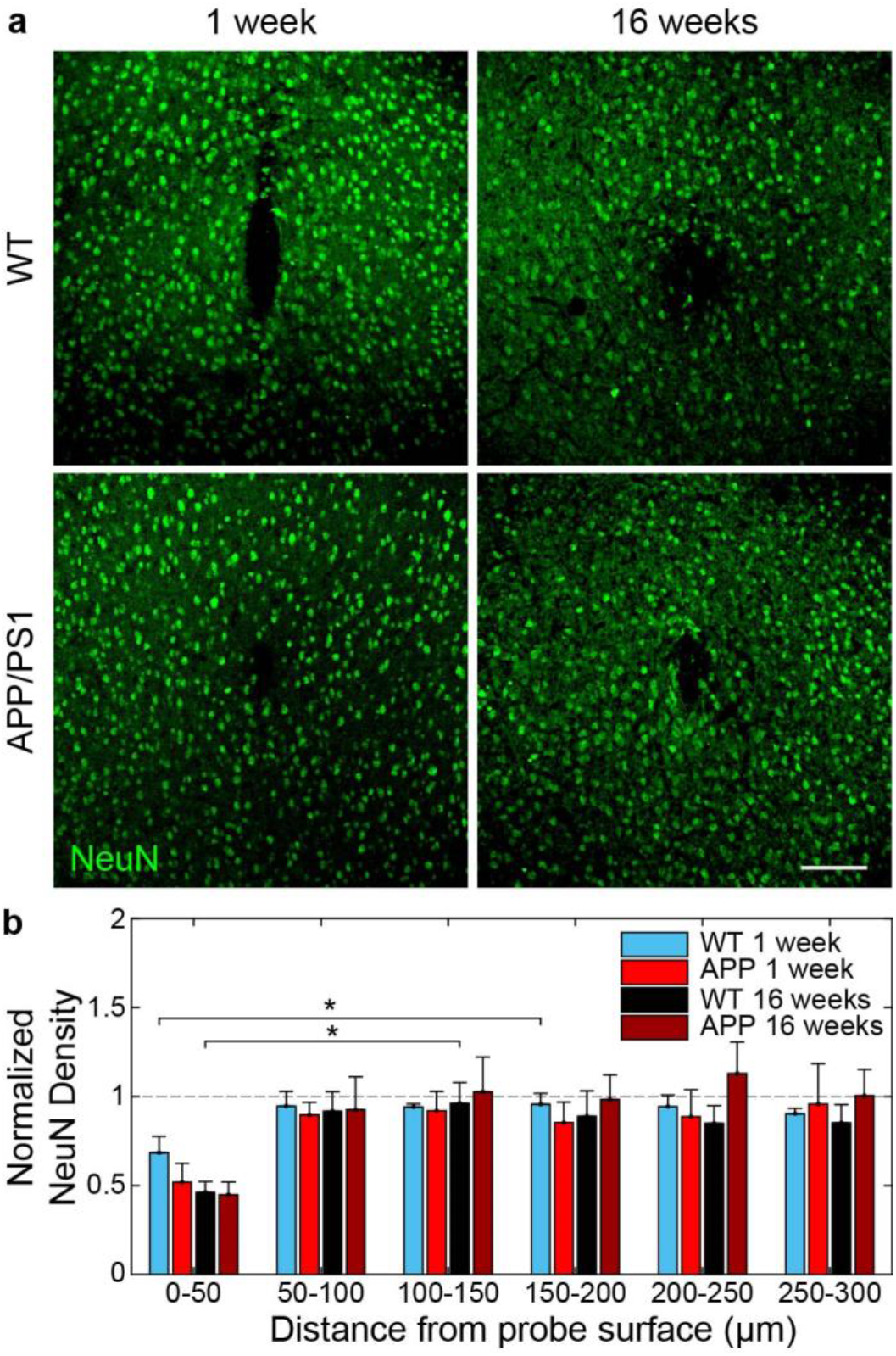
Reduced neuronal densities near chronically implanted microelectrodes. (a) Histological representation of neurons (NeuN, *green*) around chronically implanted microelectrodes in WT and APP/PS1 mice at 1- and 16-weeks post-implantation. Scale bar = 100 μm. (b) Average NeuN density within 50 μm bins up to 300 μm around chronically implanted microelectrodes at 1- and 16-weeks post-implantation in WT and APP/PS1 mice (*n* = 6 mice per group at 1 week, *n* = 7 mice per group at 16 weeks). * *p* < 0.05. All data is reported as mean ± SEM.

### 4.5. Abnormally phosphorylated tau marks regions of axonal and myelin loss around chronically implanted electrodes

Previously, we demonstrated that chronic electrode implantation leads to the re-organization of axonal and myelin fibers near the site of probe implantation^18,30^. In this study, we aimed to better understand the pathology governing axonal and myelin loss during device implantation injury and whether there is an impact following microelectrode implantation in a mouse model of AD. Staining explanted brain tissue with NF200, an axonal protein, and MBP, for myelin basic protein, we reveal unevenly distributed areas of axonal and myelin loss with close proximity to the site of microelectrode implantation (**Fig. 8a**). Interestingly, when we co-stain for AT8, a marker commonly used to study hyperphosphorylated tau in neurodegenerative disease^31^, we reveal that these areas of axon and myelin loss (i.e. NF200- and MBP-signal) correspond with abnormally high levels of AT8+ phosphorylated tau. AT8 staining intensity is visually increased near the site of probe implantation (**Fig. 8a**). Normalized fluorescence intensities of AT8 are also increased above baseline levels near the site of electrode implantation in APP/PS1 mice, but not WT mice, at 1- and 16-weeks post-implantation (**Fig. 8b**). The average normalized AT8 fluorescence intensity was significantly increased in APP/PS1 mice at 1-week post-implantation compared to WT mice, but not at 16 weeks post-implantation (**Fig. 8c,** *p*<0.01, Welch’s t-test). These results suggest that axonal and myelin pathology following device implantation injury is associated with abnormal phosphorylation of tau.

**Figure 8.**
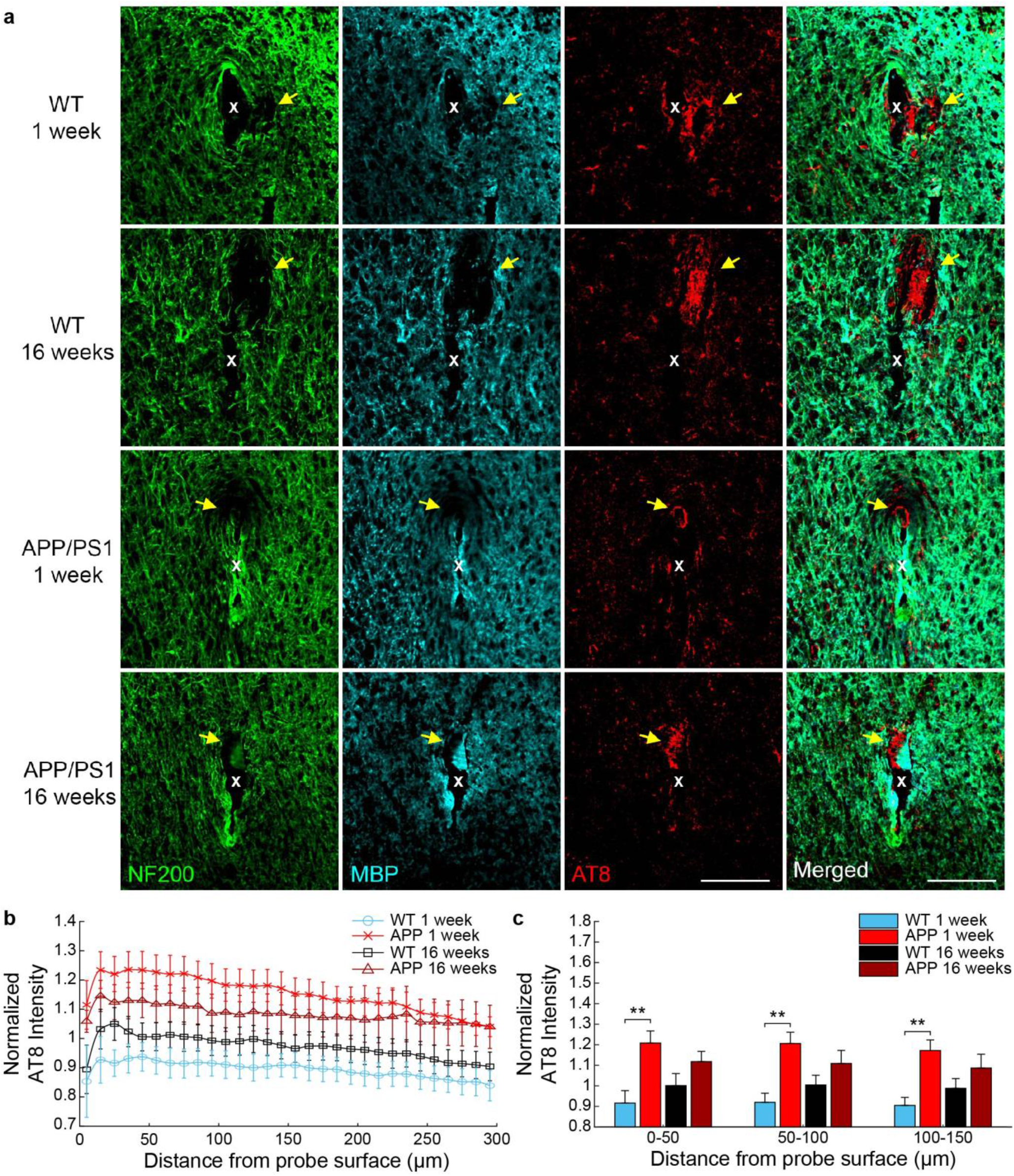
Axon and myelin pathology is associated with abnormal tau phosphorylation around chronically implanted microelectrodes. (a) Immunostaining for axons (NF200, *green*), myelin (MBP, *cyan*), and phospho-tau (AT8, *red*) reveals abnormal tau phosphorylation in areas of axon and myelin loss (*yellow arrows*) around 1- and 16-week implanted probes (white ‘x’) in WT and APP/PS1 mice. Scaler bar = 100 μm. (b) Normalized AT8 fluorescence intensity with respect to distance around chronically implanted microelectrodes at 1- and 16-weeks post-implantation in WT and APP/PS1 mice. (c) Average AT8 fluorescence intensity within 50 μm bins up to 150 μm around chronically implanted microelectrodes at 1- and 16-weeks post-implantation in WT and APP/PS1 mice (*n* = 6 mice per group at 1 week, *n* = 7 mice per group at 16 weeks). ** *p* < 0.01. All data is reported as mean ± SEM.

## 5.0. DISCUSSION

The aim of this study was to understand whether the brain’s immune response to neural electrode technology mimics neuropathology typically observed with neurodegenerative disease. Neuroinflammation, glial activation, and vascular dysfunction are all characteristic tissue events shared between both chronic implants and degenerative brain diseases, such as Alzheimer’s disease. The neuropathology of Alzheimer’s disease has been studied for more than a century, much longer than that of the foreign body response to neural microelectrodes. As a result, common biomarkers classically observed in disease, such as amyloid beta and neurofibrillary tau tangles, have been identified and implicated in the development of neurodegeneration and chronic inflammation within diseased brains. Therefore, it would be of interest to determine whether any of the identified hallmarks of neurodegeneration in Alzheimer’s disease are also involved in the biological failure mode of chronically implanted microelectrodes within the brain. Identifying predominant cellular and tissue mechanisms compromising neural health and neurotransmission around recording and stimulating brain implants will aide in the discovery and innovation of therapeutic approaches to improve the fidelity and longevity of neural electrode interfaces.

### 5.1. Accumulation of age-related lipofuscin granules following chronic electrode implantation

With aging and cellular senescence comes a natural decline in cellular metabolism, leading to the buildup of harmful metabolic waste. Lipofuscin, an auto-fluorescent composite of proteins, lipids, and sugars, accumulates naturally within the brain with age^32^. These lipofuscin granules are primarily found within lysosomes of post-mitotic and senescent cells, suggesting they are unable to be degraded normally through cellular metabolism^33,34^.. It is believed that this loss of lysosomal hydrolysis is due to decreases in lysosomal acidity and metabolic supply to sustain lysosome pH^27–29^. We report here that chronically implanted microelectrodes promote lipofuscin aggregation within the aged brain (**Fig. 2**).

Lipofuscin accumulation around microelectrodes occurred in both WT and AD mice alike, although there was an increased trend for AD mice to produce lipofuscin deposits that were greater in both number and size within the electrode microenvironment. An impairment in the clearance and removal of pathological proteins and cellular waste is a commonly proposed theory of neurodegeneration within both natural aging and brain disease. The functional role of lipofuscin deposits within the brain and whether they are neurotoxic remain poorly understood. However, lipofuscin accumulation has been previously associated with cellular oxidative stress as well as impairments in memory function, suggesting they have some impact on the health of neuronal tissue and regulation of brain activity^35^. Nevertheless, elevated accumulation of lipofuscin granules observed here around chronically implanted microelectrodes suggests that electrode implantation injury induces a backstop in the clearance mechanisms regulating removal of cellular debris and metabolic waste from the brain.

### 5.2. Patterns of amyloid deposition following chronic electrode implantation

Our initial hypothesis was that chronic microelectrode implantation would exacerbate the growth of pre-existing amyloid plaques either due to elevated deposition of neuronal or glial Aβ or due to an impairment in glial clearance of amyloid. Contrary to this hypothesis, we observed that Aβ plaques existing prior to the start of electrode implantation were, on average, smaller in volume over a 12-week implantation period compared to plaques located further away in distal brain regions (**Fig. 3**). This was the consequence of an overall negative change in size of plaques around microelectrodes. In contrast, plaques on the contralateral, uninjured hemisphere gradually increased in size due to the natural progression of neuropathology within this mouse model.

One possible explanation is that an increase in plaque size on the ipsilateral hemisphere was impaired due to a degradation in chronic imaging quality through an optical window around implanted microelectrodes over time, which would present a confound in our findings. However, in support of our two-photon findings, we observed a similar decrease in the overall area of plaques via post-mortem immunohistochemical staining confirming that the microenvironment around chronically implanted microelectrodes halts the rate of plaque growth **(Fig. 3d,e)**. It is well understood that glial cells are responsible for the degradation and clearance of amyloid from the brain and that their physiological dysfunction in Alzheimer’s disease could lead to the aberrant accumulation of amyloid, eventually forming senile plaques^36,37^. One possible interpretation of findings could be that an enhanced glial activity from the foreign body response results in the reduction in growth of nearby plaques. We show here that both microglia and astrocyte activity is upregulated with respect to chronically implanted microelectrodes in both WT and APP/PS1 mice alike **(Fig. 5–6)**.

Despite our results demonstrating that microelectrodes stall the growth of Aβ plaques locally, we also report novel amyloid deposition proximal to the electrode-tissue interface (**Fig. 4**). The amyloid clusters we observed were smaller in size compared to traditionally measured Aβ plaques yet accumulated in greater numbers throughout the course of implantation. The morphology of these clusters also appeared rounder, denser, and more symmetrical in shape, different than that of plaques which are commonly asymmetrical and consist of a dense plaque core surrounded by diffuse amyloid fibrils^31^. Microglia are known to form a neuroprotective barrier around amyloid plaques, and denser, more compact plaques are generally found to have more microglia contact^38^. One interpretation is that these amyloid clusters appear more compacted near the electrode where there is also an elevated microglial immune response and therefore more microglia association with amyloid. Alternatively, brain injury is known to increase the rate of APP processing, production of amyloid, and aggregation of Aβ plaques^39,40^ and microglia naturally phagocytose and sequester amyloid for proper degradation and removal from the brain^36^. Therefore, another possible explanation could be that these amyloid clusters do not represent typical extracellular Aβ plaque deposits but represent accumulation intracellularly within neurons and glial cells following implantation injury from chronic microelectrodes. Future studies discerning the origin of these pathological amyloid by-products could reveal additional insight on the ongoing neurodegenerative processes around neural microelectrodes.

### 5.3. Glial basis for phagocytosis and generation of amyloid precursor protein following electrode implantation

Microglia are the main potentiators of neuroinflammation both around chronic brain implants and with neurodegenerative disease. We report here that microglia activation is increased around chronically implanted microelectrodes in both WT and AD mice (**Fig. 5**). Also, we demonstrate that microglia close to the site of electrode implantation exhibit increased expression of Triggering receptor expressed on myeloid cells 2 (TREM2). TREM2 is an important metabolic receptor expressed predominantly within microglia and assists with critical microglial functions such as phagocytosis and removal of pathological waste. TREM2 has been identified as a significant risk factor in the development of Alzheimer’s disease. It is suggested that dysfunction in this critical immune cell receptor is what impairs the ability for microglia to remove amyloid and tau from Alzheimer’s brains^31^.

We demonstrate here that lipofuscin waste accumulates around chronically implanted microelectrodes in both WT and AD mice, but that AD mice trend more negatively (i.e. lipofuscin appears larger nearer to the electrode). Indeed, the hAbeta/APOE4/Trem2*R47H mouse model used in this study presents a mutation in the *Trem2* gene, potentially suggesting that increased lipofuscin accumulation around chronic microelectrodes within these mice is the result of impaired TREM2 function. Furthermore, impairment in TREM2 function interferes with the ability for microglia to form a protective barrier around amyloid plaques leading to increases in amyloid deposition and an increase in neuronal and axonal dystrophy^31^. It is also worth noting that microglia also have impaired ability to form a protective barrier around APOE4 used in this study compared to APOE3^41^. APOE4 is a major genetic risk factor for AD and is implicated in multiple neurodegenerative processes within the brain, such as amyloid/tau accumulation, neuroinflammation, impaired synaptic plasticity, vascular dysfunction, and metabolism^42^. It is unclear from the results reported here whether TREM2 or APOE function is compromised around chronically implanted microelectrodes or within AD brains. Future studies would benefit from examining the functional purpose of TREM2 and APOE within microglia at the electrode-tissue interface and how their upregulation of this phagocytic receptor contributes to the foreign body response to chronically implanted microelectrodes.

Astrocytes mediate the formation of a neurochemically impermeable and neurotoxic glial scar as part of the foreign body response to chronic brain implants. Previously, we presented a dynamic spatiotemporal pattern of astrocyte reactivity within the first few weeks during implantation using two-photon imaging around implanted microelectrodes^10^. Here, we demonstrate that astrocyte reactivity significantly increases from 1- to 16-weeks post-implantation in both WT and AD mice (**Fig. 6**). Additionally, we report astrocyte-specific upregulation of APP in proximity to chronically implanted microelectrodes within WT mice. Based on these results, it is possible that the Increase in amyloid deposition observed around microelectrode implants could be partially due to Aβ generation from reactive astrocytes. Amyloid precursor protein is a transmembrane protein expressed on cellular members and cleaved by secretases. With proper cleavage, the amyloid derived from APP does not aggregate into Aβ plaques and can be easily removed from the brain. Improper cleavage of APP, such as with–secretases, produces an aggregated form of Aβ resulting in the pathological formation of senile plaques. Future research should be performed assessing the function role of APP expression within reactive astrocytes around chronically implanted microelectrodes.

### 5.4. Neuronal and axonal pathology following chronic electrode implantation

The presence of nearby healthy neurons is critical to the long-term performance of neural microelectrode arrays. Neural densities are characteristically reduced both acute and chronically with close spatial proximity to implanted microelectrodes^43–45^. In this study, we confirm that microelectrodes impact the number of neurons located adjacent to the site of implantation (**Fig. 7**). Despite a visible difference in the density of neurons within the 0-50 μm tissue region around chronically implanted microelectrodes at 1- and 16-weeks post-implantation, we did not detect any significant differences between WT and APP/PS1 mice. It is important to keep in mind that at 16-weeks post-implantation APP/PS1 mice are at 6 months of age and neuronal numbers are not expected to be naturally altered within the AD model at this age^46–48^. Furthermore, our analyses are only limited to understanding whether NeuN+ neurons are present or not around the electrode compared to intact control regions but does not discern whether the neurons that remain are viable or whether they may be otherwise functionally impaired around chronically implanted devices. In support of this, there have been reports of explanted tissue populated by many neurons yet still demonstrating poor electrophysiological recording performance^5,49^. One possible explanation is that the neurons which were not directly impacted by implantation injury are quiescent or their activity is somehow suppressed^5^. It is also important to note that trauma to the brain can alter the expression of neuronal proteins traditionally used to evaluate neuronal health and viability, such as NeuN, in the absence of any apparent neurodegeneration^50^. Therefore, further *in vivo* evaluation is needed to fully understand the physiological consequences imparted on neurons around chronically implanted microelectrodes and in rodent models of neurodegenerative disease.

Maintaining axonal and myelin integrity around chronically implanted electrodes is essential for effective transmission of neuronal information and excitability of neurons in brain regions both near and far from intracortical recording and stimulating electrodes. We have previously demonstrated re-organization and potential sprouting of axons as well as spatiotemporal patterns of axonal and myelin blebbing around intracortical electrodes^5,7,8^. Here, we demonstrate that electrode implantation can induce regions of axonal and myelin loss with proximity to the site of electrode insertion (**Fig. 8**). A previous study investigating local markers of tissue inflammation around chronically implanted microelectrodes noted the appearance of hyperphosphorylated tau at the lesion border within explanted brain tissue following 16-weeks of electrode implantation in rats^20^. This same study also noted similar neuropathology around the site of device insertion following 5 months of implantation in a human Parkinsonian patient, suggesting this phenomenon can occur in both rodents and humans. In line with this study, we observe elevated expression of phosphorylated tau near chronically implanted microelectrodes at 1- and 16-weeks post-implantation in both WT and AD mice. To our surprise, these areas of increased tau phosphorylation overlap with identified tissue regions lacking axons and myelin, suggesting that abnormal tau phosphorylation is an indicator of axonal and myelin pathology around chronically implanted microelectrodes. Phosphorylation of tau is a naturally occurring phenomenon important for microtubule assembly, axonal transport, and neuronal plasticity^51–53^. Hyperphosphorylated tau, however, is a pathological hallmark of Alzheimer’s disease in which misfolded tau proteins aggregate and form neurofibrillary tau tangles within axons, interfering with neuronal information transmission^53^. It is unclear from these findings whether hyperphosphorylated tau is the symptom or cause of axonal and myelin pathology around chronically implanted microelectrodes or whether abnormal tau phosphorylation would have a significant impact on the recording or excitability of neurons at the electrode-tissue interface. Understanding if tau phosphorylation precedes axonal and myelin loss (or vice versa) and if tau phosphorylation contributes to functional electrode performances could point towards novel biological targets to address neurodegeneration around chronic brain implants.

### 5.5. Future directions

We determined that the levels of lipofuscin, APP, and Aβ are all upregulated around chronically implanted microelectrodes. While the pathological consequence of lipofuscin remains a mystery, it demonstrates no useful physiological purpose to-date. Previous work has demonstrated that chronic treatment with the antioxidant melatonin can reduce lipofuscin content within the rat hippocampus^54^. Melatonin has previously shown to improve both microglia activation and recording performance of chronically implanted electrodes^55,56^, suggesting targeting glial function in general could be an effective way to address lipofuscin aggregation indirectly. APP itself is important for neural stem cell proliferation and axon outgrowth following injury and therefore may be neuroprotective in certain pathological conditions^57,58^. However, impaired cleavage of APP by β-secretases generates precursors for Aβ deposition. It is currently unclear how the levels of β-secretase, as well as α- and γ-secretase, are impacted around chronically implanted microelectrodes. Finally, previous and current FDA-approved clinical trials have focused on improving cognitive and behavioral outcomes in Alzheimer’s disease by using monoclonal antibodies to target and promote clearance of Aβ plaques from the brain, yet the results have either been mixed or unsuccessful thus far^14,59^.

There is current debate on whether the presence of amyloid is even a cause or just a symptom of neurodegeneration within Alzheimer’s disease (i.e. “amyloidogenic” vs. “non-amyloidogenic” hypothesis)^60^. In any case, the abnormal accumulation of lipofuscin, APP, and amyloid around chronically implanted electrodes suggests a critical failure in tissue clearance mechanisms and may hint towards the accumulation of other potentially harmful factors, such as reactive oxidative species and misfolded tau proteins, whose improper removal from the brain can be detrimental to neural health and function. Future studies should determine what other potential tissue factors accumulating at the electrode-tissue interface to develop a more holistic understanding of the potentially detrimental biochemical processes in effect around chronically implanted electrodes.

Tau phosphorylation is mediated by a number of different kinases, such as cyclin-dependent kinase 5 (cdk5) and glycogen synthase kinase-3 (GSK-3)^61,62^, and phosphatases, such as protein phosphatase PP1, PP2A, and PP5^63^. Accounting for the changes in expression levels of these critical regulators of tau phosphorylation around microelectrodes could point toward novel targets during neurodegeneration surrounding chronic electrode implantation. There is also evidence that the presence of amyloid can alter tau phosphorylation^64,65^. It is important to note that the methoxy-X04 label used in this study binds to both amyloid and tau within the brain^66,67^. Therefore, it is unclear whether the methoxy-X04 labeling around implanted microelectrodes is completely representative of amyloid protein or if there is also partial labeling of tau as well. Phosphorylated tau was also reported here as being increased in unusual densities around chronically implanted microelectrodes. Future research should use more targeted investigative methods, such as using P130S mouse models more directed at studying tau pathology in AD, to disentangle the potential relationship between amyloid and tau at the electrode-tissue interface.

Overall, we did not report much significance in the difference between WT and AD mice for some of the patterns of neuropathology commonly shared between the two models. For two-photon imaging experiments, mice were aged to 6 months prior to electrode implantation to understand the impact of electrode implantation injury on the rate of growth of nearby pre-existing Aβ plaques. For histology experiments, APP/PS1 were implanted at 2 months of age and, by 16-weeks of implantation, mice were only 6 months old. At this age, APP/PS1 mice are just beginning to present amyloid deposition and therefore it is still relatively early in the progression of AD pathology typical for this mouse model. It could be, when comparing AD mice of a year or more in age to similarly age-matched controls, that we may observe a much greater difference in brain tissue responses following chronic microelectrode implantation. Despite this, we were still able to show that microelectrode implantation accelerates the onset of amyloid pathology even within 2-month-old APP/PS1 mice. Additionally, we reported novel findings on the expression of various cell and tissue factors related to aging and neurodegenerative disease around implanted microelectrodes which will aide future investigators in narrowing their research focuses and develop more informed intervention strategies for addressing the foreign body reaction to chronic brain implants.

## 6.0. CONCLUSION

The power of neural interface technology to aide in the discovery of previously unknown neuroscientific phenomenon and provide means for effective clinical therapy of neurological dysfunction is limited due to yet unresolved tissue responses to implanted recording and stimulating electrodes within the brain. These presently elusive biological mechanisms impairing the long-term application of neural microelectrodes could potentially be revealed by referencing the study of other brain disorders similarly plagued by neuronal, glial, and vascular dysfunction, such as Alzheimer’s disease. In this study, we employed mouse models of Alzheimer’s disease to reveal the change in expression of various tissue factors that offer new insights on neurodegeneration surrounding chronic brain implants. We determined that microelectrode implantation preferentially favors the accumulation of pathological proteins related to aging and neurological disease at the electrode-tissue interface. We also highlighted novel spatiotemporal patterns of glial-specific expression of different factors related to deposition and clearance of amyloid within the brain as well as new information on potential neurodegenerative mechanisms governing axonal pathology around chronic microelectrodes. In summary, these research findings provide a new perspective regarding the progression of neurodegenerative injury surrounding neural interface technology and will aide in more rigorous study of electrode-tissue biocompatibility as well as expedited development of advanced therapies for both brain injury and disease.

## 7.0. ACKNOWLEDGEMENTS

The authors would like to thank Dr. Amantha Thathiah, Thais Guimaraes, and Keying Chen for consultation on immunohistology experiments. This work was supported by NIH NINDS R21NS108098, NIH NINDS R01NS094396 and a diversity supplement to the parent grant, NIH NIA R03AG072218, and NIH NINDS F99NS124186.

## REFERENCES

1 Schwartz, A. B., Cui, X. T., Weber, D. J. & Moran, D. W. Brain-controlled interfaces: movement restoration with neural prosthetics. Neuron 52, 205–220, doi:10.1016/j.neuron.2006.09.019 (2006).

2 Flesher, S. N. et al. Intracortical microstimulation of human somatosensory cortex. Science Translational Medicine 8, 361ra141–361ra141, doi:doi:10.1126/scitranslmed.aaf8083 (2016).

3 Hughes, C. L. et al. Perception of microstimulation frequency in human somatosensory cortex. Elife 10, e65128 (2021).

4 Kozai, T. D., Jaquins-Gerstl, A. S., Vazquez, A. L., Michael, A. C. & Cui, X. T. Brain tissue responses to neural implants impact signal sensitivity and intervention strategies. ACS chemical neuroscience 6, 48–67 (2015).

5 Michelson, N. J. et al. Multi-scale, multi-modal analysis uncovers complex relationship at the brain tissue-implant neural interface: new emphasis on the biological interface. Journal of neural engineering 15, 033001 (2018).

6 Wellman, S. M. & Kozai, T. D. Vol. 8 2578–2582 (ACS Publications, 2017).

7 Wellman, S. M., Li, L., Yaxiaer, Y., McNamara, I. & Kozai, T. D. Revealing spatial and temporal patterns of cell death, glial proliferation, and blood-brain barrier dysfunction around implanted intracortical neural interfaces. Frontiers in neuroscience 13, 493 (2019).

8 Chen, K., Wellman, S. M., Yaxiaer, Y., Eles, J. R. & Kozai, T. D. In vivo spatiotemporal patterns of oligodendrocyte and myelin damage at the neural electrode interface. Biomaterials 268, 120526 (2021).

9 Kozai, T. D. Y. et al. Reduction of neurovascular damage resulting from microelectrode insertion into the cerebral cortex using in vivo two-photon mapping. Journal of neural engineering 7, 046011 (2010).

10 Savya, S. P. et al. In vivo spatiotemporal dynamics of astrocyte reactivity following neural electrode implantation. Biomaterials 289, 121784 (2022).

11 Hughes, C. L. et al. Neural stimulation and recording performance in human sensorimotor cortex over 1500 days. Journal of Neural Engineering 18, 045012 (2021).

12 Vickers, J. C. et al. The cause of neuronal degeneration in Alzheimer’s disease. Progress in neurobiology 60, 139–165 (2000).

13 Zlokovic, B. V. Neurovascular pathways to neurodegeneration in Alzheimer’s disease and other disorders. Nature Reviews Neuroscience 12, 723–738 (2011).

14 Salloway, S. et al. Two phase 3 trials of bapineuzumab in mild-to-moderate Alzheimer’s disease. New England Journal of Medicine 370, 322–333 (2014).

15 Guerreiro, R. et al. TREM2 variants in Alzheimer’s disease. New England Journal of Medicine 368, 117–127 (2013).

16 Neu, S. C. et al. Apolipoprotein E genotype and sex risk factors for Alzheimer disease: a meta-analysis. JAMA neurology 74, 1178–1189 (2017).

17 Sims, R. et al. Rare coding variants in PLCG2, ABI3, and TREM2 implicate microglial-mediated innate immunity in Alzheimer’s disease. Nature genetics 49, 1373–1384 (2017).

18 Cations, M. et al. Non-Genetic Risk Factors for Degenerative and Vascular Young Onset Dementia: Results from the INSPIRED and KGOW Studies. Journal of Alzheimer’s disease: JAD 62, 1747–1758, doi:10.3233/jad-171027 (2018).

19 Gardner, R. C. et al. Dementia risk after traumatic brain injury vs nonbrain trauma: the role of age and severity. JAMA neurology 71, 1490–1497 (2014).

20 McConnell, G. C. et al. Implanted neural electrodes cause chronic, local inflammation that is correlated with local neurodegeneration. Journal of neural engineering 6, 056003 (2009).

21 Kozai, T. D. Y., Vazquez, A. L., Weaver, C. L., Kim, S.-G. & Cui, X. T. In vivo two-photon microscopy reveals immediate microglial reaction to implantation of microelectrode through extension of processes. Journal of neural engineering 9, 066001 (2012).

22 Wellman, S. M. & Kozai, T. D. In vivo spatiotemporal dynamics of NG2 glia activity caused by neural electrode implantation. Biomaterials 164, 121–133 (2018).

23 Kozai, T. D., Eles, J. R., Vazquez, A. L. & Cui, X. T. Two-photon imaging of chronically implanted neural electrodes: Sealing methods and new insights. Journal of neuroscience methods 258, 46–55 (2016).

24 Kozai, T. D. et al. Chronic tissue response to carboxymethyl cellulose based dissolvable insertion needle for ultra-small neural probes. Biomaterials 35, 9255–9268 (2014).

25 Elder, G. A., Gama Sosa, M. A. & De Gasperi, R. Transgenic Mouse Models of Alzheimer’s Disease. Mount Sinai Journal of Medicine: A Journal of Translational and Personalized Medicine 77, 69–81, doi:https://doi.org/10.1002/msj.20159 (2010).

26 Yan, P. et al. Characterizing the appearance and growth of amyloid plaques in APP/PS1 mice. The Journal of neuroscience: the official journal of the Society for Neuroscience 29, 10706–10714, doi:10.1523/jneurosci.2637-09.2009 (2009).

27 Stoka, V., Turk, V. & Turk, B. Lysosomal cathepsins and their regulation in aging and neurodegeneration. Ageing research reviews 32, 22–37 (2016).

28 Guha, S., Liu, J., Baltazar, G., Laties, A. M. & Mitchell, C. H. Rescue of compromised lysosomes enhances degradation of photoreceptor outer segments and reduces lipofuscin-like autofluorescence in retinal pigmented epithelial cells. Retinal Degenerative Diseases, 105–111 (2014).

29 Kurz, D. J., Decary, S., Hong, Y. & Erusalimsky, J. D. Senescence-associated (beta)-galactosidase reflects an increase in lysosomal mass during replicative ageing of human endothelial cells. Journal of cell science 113, 3613–3622 (2000).

30 D’Andrea, M. R., Cole, G. M. & Ard, M. D. The microglial phagocytic role with specific plaque types in the Alzheimer disease brain. Neurobiology of aging 25, 675–683 (2004).

31 Yuan, P. et al. TREM2 Haplodeficiency in Mice and Humans Impairs the Microglia Barrier Function Leading to Decreased Amyloid Compaction and Severe Axonal Dystrophy. Neuron 90, 724–739, doi:10.1016/j.neuron.2016.05.003 (2016).

32 Gray, D. A. & Woulfe, J. Lipofuscin and Aging: A Matter of Toxic Waste. Science of Aging Knowledge Environment 2005, re1-re1, doi:doi:10.1126/sageke.2005.5.re1 (2005).

33 Porta, E. A. Pigments in Aging: An Overview. Annals of the New York Academy of Sciences 959, 57–65, doi:https://doi.org/10.1111/j.1749-6632.2002.tb02083.x (2002).

34 Oxidative Stress, Accumulation of Biological ‘Garbage’, and Aging. Antioxidants & Redox Signaling 8, 197–204, doi:10.1089/ars.2006.8.197 (2006).

35 Keller, J. N. et al. Autophagy, proteasomes, lipofuscin, and oxidative stress in the aging brain. The international journal of biochemistry & cell biology 36, 2376–2391 (2004).

36 Lai, A. Y. & McLaurin, J. Clearance of amyloid-β peptides by microglia and macrophages: the issue of what, when and where. Future neurology 7, 165–176, doi:10.2217/fnl.12.6 (2012).

37 Lee, C. & Landreth, G. E. The role of microglia in amyloid clearance from the AD brain. Journal of neural transmission 117, 949–960 (2010).

38 Condello, C., Yuan, P., Schain, A. & Grutzendler, J. Microglia constitute a barrier that prevents neurotoxic protofibrillar Aβ42 hotspots around plaques. Nature communications 6, 6176, doi:10.1038/ncomms7176 (2015).

39 Johnson, V. E., Stewart, W. & Smith, D. H. Traumatic brain injury and amyloid-β pathology: a link to Alzheimer’s disease? Nature reviews. Neuroscience 11, 361–370, doi:10.1038/nrn2808 (2010).

40 Washington, P. M., Morffy, N., Parsadanian, M., Zapple, D. N. & Burns, M. P. Experimental traumatic brain injury induces rapid aggregation and oligomerization of amyloid-beta in an Alzheimer’s disease mouse model. Journal of neurotrauma 31, 125–134, doi:10.1089/neu.2013.3017 (2014).

41 Fitz, N. F. et al. Phospholipids of APOE lipoproteins activate microglia in an isoform-specific manner in preclinical models of Alzheimer’s disease. Nature communications 12, 3416 (2021).

42 Safieh, M., Korczyn, A. D. & Michaelson, D. M. ApoE4: an emerging therapeutic target for Alzheimer’s disease. BMC medicine 17, 1–17 (2019).

43 Shoffstall, A. J. et al. Characterization of the neuroinflammatory response to thiol-ene shape memory polymer coated intracortical microelectrodes. Micromachines 9, 486 (2018).

44 McCreery, D., Cogan, S., Kane, S. & Pikov, V. Correlations between histology and neuronal activity recorded by microelectrodes implanted chronically in the cerebral cortex. Journal of neural engineering 13, 036012 (2016).

45 Potter-Baker, K. A. et al. A comparison of neuroinflammation to implanted microelectrodes in rat and mouse models. Biomaterials 35, 5637–5646 (2014).

46 Szögi, T. et al. Examination of Longitudinal Alterations in Alzheimer’s Disease-Related Neurogenesis in an APP/PS1 Transgenic Mouse Model, and the Effects of P33, a Putative Neuroprotective Agent Thereon. International journal of molecular sciences 23, 10364 (2022).

47 Liu, L. et al. Multiple inflammatory profiles of microglia and altered neuroimages in APP/PS1 transgenic AD mice. Brain research bulletin 156, 86–104 (2020).

48 Zhang, J. et al. Long-term treadmill exercise attenuates Aβ burdens and astrocyte activation in APP/PS1 mouse model of Alzheimer’s disease. Neuroscience letters 666, 70–77 (2018).

49 Kozai, T. D. et al. Effects of caspase-1 knockout on chronic neural recording quality and longevity: insight into cellular and molecular mechanisms of the reactive tissue response. Biomaterials 35, 9620–9634 (2014).

50 Munoz-Ballester, C., Mahmutovic, D., Rafiqzad, Y., Korot, A. & Robel, S. Mild Traumatic Brain Injury-Induced Disruption of the Blood-Brain Barrier Triggers an Atypical Neuronal Response. Frontiers in cellular neuroscience 16, 32 (2022).

51 Dixit, R., Ross, J. L., Goldman, Y. E. & Holzbaur, E. L. Differential regulation of dynein and kinesin motor proteins by tau. Science 319, 1086–1089 (2008).

52 Barbier, P. et al. Role of tau as a microtubule-associated protein: structural and functional aspects. Frontiers in aging neuroscience 11, 204 (2019).

53 Noble, W., Hanger, D. P., Miller, C. C. & Lovestone, S. The importance of tau phosphorylation for neurodegenerative diseases. Frontiers in neurology 4, 83 (2013).

54 Abd El Mohsen, M. M. et al. Age-associated changes in protein oxidation and proteasome activities in rat brain: modulation by antioxidants. Biochemical and biophysical research communications 336, 386–391 (2005).

55 Krahe, D. D., Woeppel, K. M., Yang, Q., Kushwah, N. & Cui, X. T. Melatonin Decreases Acute Inflammatory Response to Neural Probe Insertion. Antioxidants 11, 1628 (2022).

56 Golabchi, A. et al. Melatonin improves quality and longevity of chronic neural recording. Biomaterials 180, 225–239 (2018).

57 Caillé, I. et al. Soluble form of amyloid precursor protein regulates proliferation of progenitors in the adult subventricular zone. (2004).

58 Wang, X., Huang, T., Bu, G. & Xu, H. Dysregulation of protein trafficking in neurodegeneration. Molecular neurodegeneration 9, 1–9 (2014).

59 Honig, L. S. et al. Trial of solanezumab for mild dementia due to Alzheimer’s disease. New England Journal of Medicine 378, 321–330 (2018).

60 Herrup, K. The case for rejecting the amyloid cascade hypothesis. Nature neuroscience 18, 794–799 (2015).

61 Baumann, K., Mandelkow, E.-M., Biernat, J., Piwnica-Worms, H. & Mandelkow, E. Abnormal Alzheimer-like phosphorylation of tau-protein by cyclin-dependent kinases cdk2 and cdk5. FEBS letters 336, 417–424 (1993).

62 Hanger, D. P., Hughes, K., Woodgett, J. R., Brion, J.-P. & Anderton, B. H. Glycogen synthase kinase-3 induces Alzheimer’s disease-like phosphorylation of tau: generation of paired helical filament epitopes and neuronal localisation of the kinase. Neuroscience letters 147, 58–62 (1992).

63 Liu, F., Grundke-Iqbal, I., Iqbal, K. & Gong, C. X. Contributions of protein phosphatases PP1, PP2A, PP2B and PP5 to the regulation of tau phosphorylation. European Journal of Neuroscience 22, 1942–1950 (2005).

64 Pigino, G., Pelsman, A., Mori, H. & Busciglio, J. Presenilin-1 mutations reduce cytoskeletal association, deregulate neurite growth, and potentiate neuronal dystrophy and tau phosphorylation. Journal of Neuroscience 21, 834–842 (2001).

65 Busciglio, J., Lorenzo, A., Yeh, J. & Yankner, B. A. β-Amyloid fibrils induce tau phosphorylation and loss of microtubule binding. Neuron 14, 879–888 (1995).

66 Kuchibhotla, K. V. et al. Neurofibrillary tangle-bearing neurons are functionally integrated in cortical circuits in vivo. Proceedings of the National Academy of Sciences of the United States of America 111, 510–514, doi:10.1073/pnas.1318807111 (2014).

67 Klunk, W. E. et al. Imaging Aβ Plaques in Living Transgenic Mice with Multiphoton Microscopy and Methoxy-X04, a Systemically Administered Congo Red Derivative. Journal of Neuropathology & Experimental Neurology 61, 797–805, doi:10.1093/jnen/61.9.797 (2002).

